# SHAPR predicts 3D cell shapes from 2D microscopic images

**DOI:** 10.1101/2021.09.29.462353

**Authors:** Dominik J. E. Waibel, Niklas Kiermeyer, Scott Atwell, Ario Sadafi, Matthias Meier, Carsten Marr

**Affiliations:** Institute of AI for Health, Helmholtz Munich - German Research Center for Environmental Health, Neuherberg, Germany; Institute of Computational Biology, Helmholtz Munich - German Research Center for Environmental Health, Neuherberg, Germany; Technical University of Munich, School of Life Sciences, Weihenstephan, Germany; Computer Aided Medical Procedures, Technical University of Munich, Munich, Germany; Helmholtz Pioneer Campus, Helmholtz Munich - German Research Center for Environmental Health, Neuherberg, Germany

**Keywords:** 3D shape prediction, stereology, single-cell morphometry, deep learning, autoencoder, adversarial learning

## Abstract

Reconstruction of shapes and sizes of three-dimensional (3D) objects from two-dimensional (2D) information is an intensely studied subject in computer vision. We here consider the level of single cells and nuclei and present a neural network-based SHApe PRediction autoencoder. For proof-of-concept, SHAPR reconstructs 3D shapes of red blood cells from single view 2D confocal microscopy images more accurately than naïve stereological models and significantly increases the feature-based prediction of red blood cell types from F1 = 79.0% to F1 = 87.4%. Applied to 2D images containing spheroidal aggregates of densely grown human induced pluripotent stem cells, we find that SHAPR learns fundamental shape properties of cell nuclei and allows for prediction-based morphometry. Reducing imaging time and data storage, SHAPR will help to optimize and up-scale image-based high-throughput applications for biomedicine.

## Introduction

Recording single cells in three dimensions (3D) for high-throughput biomedical applications is prohibitively time-consuming as it requires the acquisition of multiple two-dimensional (2D) images. This raises the question of how to optimally trade-off between throughput and resolution in space and time. A number of methods to reduce imaging time for single cell characterization have recently been developed, ranging from microscopic techniques, such as optical diffraction tomography (Balasubramani et al., 2021), to in-silico fluorescence staining (Christiansen et al., 2018; Ounkomol et al., 2018; Rivenson et al., 2019) and image restoration techniques (Weigert et al., 2018). To predict single cell 3D morphology, one would ideally be able to exploit the information in 2D fluorescence microscopy images. Deep learning-based solutions for predicting 3D object shapes from photographs exist, creating meshes (Gkioxari et al., 2019; Wang et al., 2018), voxels (Choy et al., 2016), or point clouds (Fan et al., 2017) for airplanes, cars, and furniture, but they cannot be translated to fluorescence microscopy for several reasons. First, fluorescence microscopy imaging is fundamentally different from real-world photographs in terms of color, contrast, and object orientation. Second, unlike the shapes of cars or furniture that might vary due to differing photographic viewpoints, the shapes of single cells are similar but never the same, and it is often not feasible to image the same cell from different angles in high throughput microscopy. Finally, existing computer vision algorithms have been trained on tens of thousands of photographs where synthetic 3D models are available (Chang et al., 2015; Sun et al., 2018; Xiang et al., 2014). In the biomedical domain the number of potential training images is orders of magnitude smaller. While Wu et al. (2019) have demonstrated that neural networks can be used for 3D refocusing onto a user-defined surface from 2D microscopy images containing single fluorescent beads or fluorophore signals, (Wu et al., 2019) to the best of our knowledge no model exists in the biomedical domain to reconstruct 3D cell shapes from 2D confocal microscopy images.

## Results

We addressed these problems with SHAPR, a deep learning network algorithm that combines a 2D encoder for feature extraction from 2D images with a 3D decoder to predict 3D shapes from a latent space representation (Fig. 1a and Supplementary Fig. 1). For proof of concept, we predicted cell shapes using a recently published library detailing 3D red blood cell shapes (n = 825 cells) (Simionato et al., 2021). Each cell shape was reconstructed from 68 confocal images with a z-resolution of 0.3μm. Using the 2D image that intersects the red blood cell at the center slice, and the corresponding segmentation as input, SHAPR was trained by minimizing binary cross-entropy and dice loss between the true and the predicted 3D red blood cell shape (Fig. 1b and Methods). To increase SHAPR’s prediction accuracy, a discriminator model (Fig. 1b) was trained to differentiate between true and predicted 3D cell shapes. SHAPR and the discriminator were trained until the predicted cell shape converged to an optimum. In each one of five cross-validation runs, 495 (60%) red blood cells from the library were used for training and 165 (20%) for intermediate validation during training. During testing, we predicted the 3D shapes of 165 (20%) previously unseen red blood cells. The results demonstrate that SHAPR is able to predict single red blood cell 3D shapes: while non-complex morphologies from red blood cells with a biconcave discoid shape (stomatocyte-discocyte-echinocyte (SDE) shape class) were predicted with low relative volume error, more complex shapes with irregular protrusions or cavities as seen in knizocytes and acanthocytes were predicted with larger errors (Fig. 1c). We compared this cell shape prediction to two naïve stereological models, i.e. a cylindrical and an ellipsoid fit (see Methods). SHAPR predictions significantly outperformed these models with respect to volume error, 2D surface area error, surface roughness error and intersection over union (Fig. 1d).

**Fig. 1.**
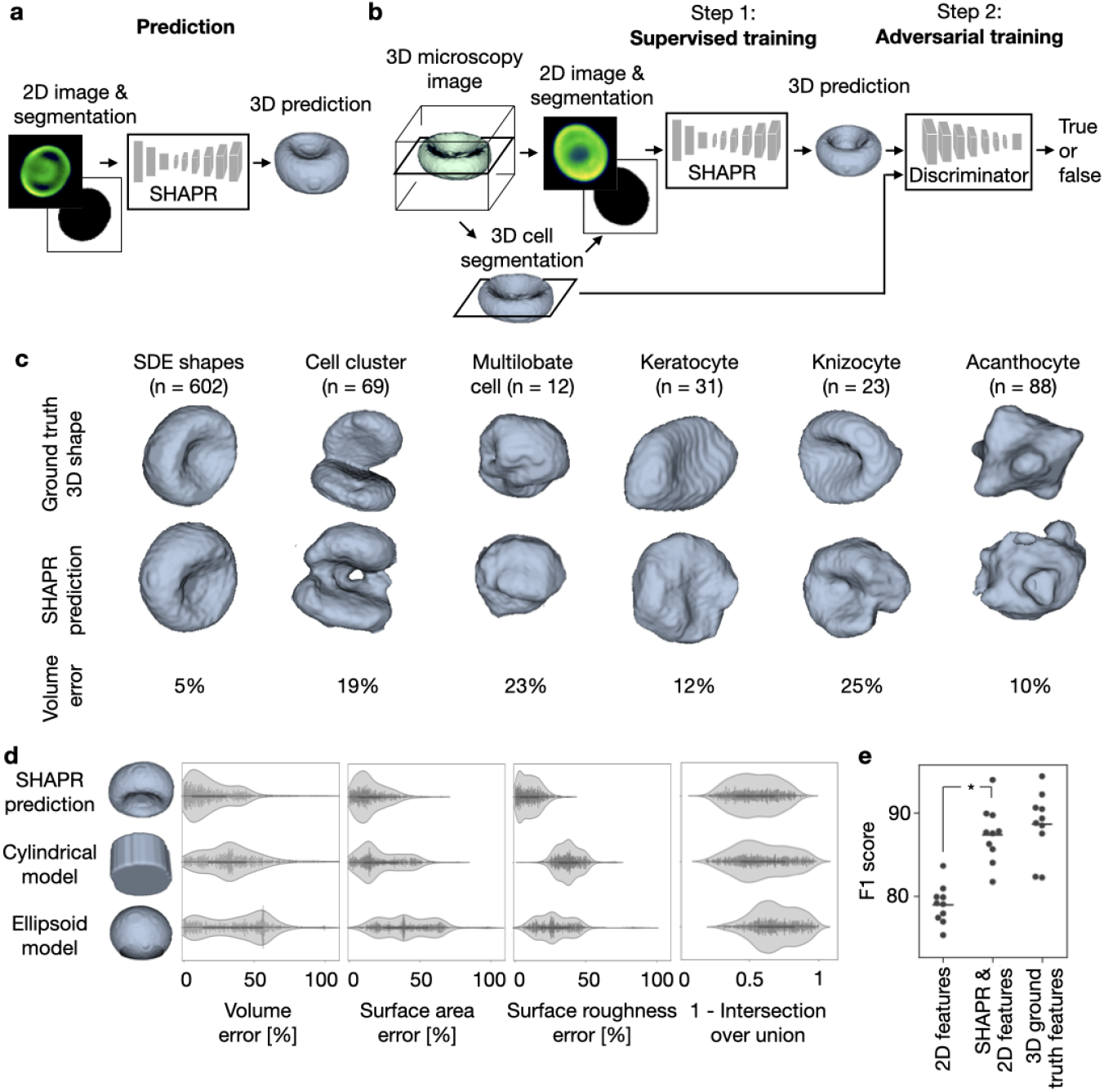
SHAPR predicts 3D cell shapes from 2D microscopic images more accurately than naïve stereological models and improves shape classification. **a**, SHAPR consists of an encoder for embedding 2D images into a 128-dimensional latent space and a decoder for reconstructing 3D cell shapes from the latent space representations. **b**, Two-step training approach: during step 1, SHAPR was trained in a supervised fashion with 2D fluorescent confocal cell microscopy images and their corresponding binary segmentations from a red blood cell library. During step 2, SHAPR was fine-tuned with a discriminator challenging its cell shape predictions. **c**, Example predictions for a set of red blood cells representing six different classes. The SDE shape class combines spherocytes, stomatocytes, discocytes, and echinocytes. **d**, The volume error is significantly lower for SHAPR (20 ± 18%) as compared to two naïve stereological models (Volume_cylindrical_ = 33 ± 22, p_cylindrical_ = 2.6 _×_ 10^−46^; and Volume_ellipsoid_ = 37 ± 23; p_ellipsoid_ = 7.8_×_10^−73^, n = 825, paired Wilcoxon signed-rank test). Volume, surface area, and roughness error are significantly reduced. **e**, Random forest-based red blood cell classification is significantly improved when morphological features extracted from SHAPR predicted cell shapes are added to features derived from 2D images (p=0.005, paired Wilcoxon signed-rank test, n = 825).

Simionato et al. (2021) classified red blood cells into six categories (Fig. 1c) based on their 3D morphology. Can SHAPR predictions from 2D images improve such a downstream classification task? To investigate this, we extracted 126 morphological features, 11 features from an additionally predicted object mesh, object moments up to third order, correlation and dissimilarity of gray level co-occurrence matrices, and 64 Gabor features (see Methods for more details) from each predicted 3D cell shape. Using random forest (Breiman, 2001), we classified each blood cell into one of the six classes and compared SHAPR’s performance with the 3D ground truth features and a 2D baseline, where only features derived from the 2D image and segmentation were used (see Methods). As expected, classification based on ground truth features led to the highest F1 score (Fig. 1e; 88.6 ± 3.7%). Strikingly, enriching 2D features with SHAPR derived features performed significantly better (F1 = 87.4 ± 3.1%, mean±std.dev., n=10 cross-validation runs) than using 2D features only (F1 = 79.0 ± 2.2%) in a tenfold cross-validation (Fig. 1e, p = 0.005, paired Wilcoxon signed-rank test).

Predicting shapes from 2D planes close to the cell’s center of mass does not accurately reflect the complexity of real world applications. Therefore, we challenged SHAPR with the task of predicting cell nuclei shapes from confocal z-stacks containing fluorescence counterstained nuclei from human-induced pluripotent stem cells (iPSCs) cultured in a spheroidal aggregate. To generate the ground truth data, 887 cell nuclei from six iPSC-derived aggregates were manually segmented in 3D (Fig. 2a). SHAPR was provided with one 2D image slice taken at an aggregate depth of 22 μm and it’s the corresponding segmentation as input (Fig. 2b). Nuclei were thus cut at random heights, leading to a variety of segmented areas, markedly complicating the prediction of 3D shapes (Fig. 2c). Following this, we trained five SHAPR models during cross-validation. Predictions were compared to cylindrical and ellipsoid fits, as described above. Again, the relative volume error was significantly lower for SHAPR (Fig. 2d; Volume_SHAPR_ = 33 ± 41 vs. Volume_Cylindrical_ = 44 ± 25, p = 9.2 _×_ 10^−36^, and Volume_Ellipsoid_ = 62 ± 29, p = 8.7 _×_ 10^−86^, n = 887, paired Wilcoxon signed-rank test) compared to the naïve models. More importantly, SHAPR predictions were also closer to the true nuclei shapes in terms of volume and surface area compared with cylindrical and ellipsoid model predictions (Fig. 2d). To determine how much information our model had learned about nuclear shape, we compared the 2D segmentation area with the volume of the ground truth, the SHAPR predictions, and the cylindrical and ellipsoid fits (Fig. 2e). As expected, the cylindrical and ellipsoid models were simply extrapolating area to volume monotonically, while the ground truth suggested a more complex relationship. SHAPR was able to learn that small segmentation areas (<200 pixels) do not emerge from minuscule nuclei but from slices at the nuclear edge. Notably, our model could only obtain high intersection over union scores for slices close to the center of mass, in contrast to volume and surface error (Fig. 2f). This suggests that it can predict volume and surface correctly, but not if a nucleus is cut in its upper or the lower half.

**Fig. 2.**
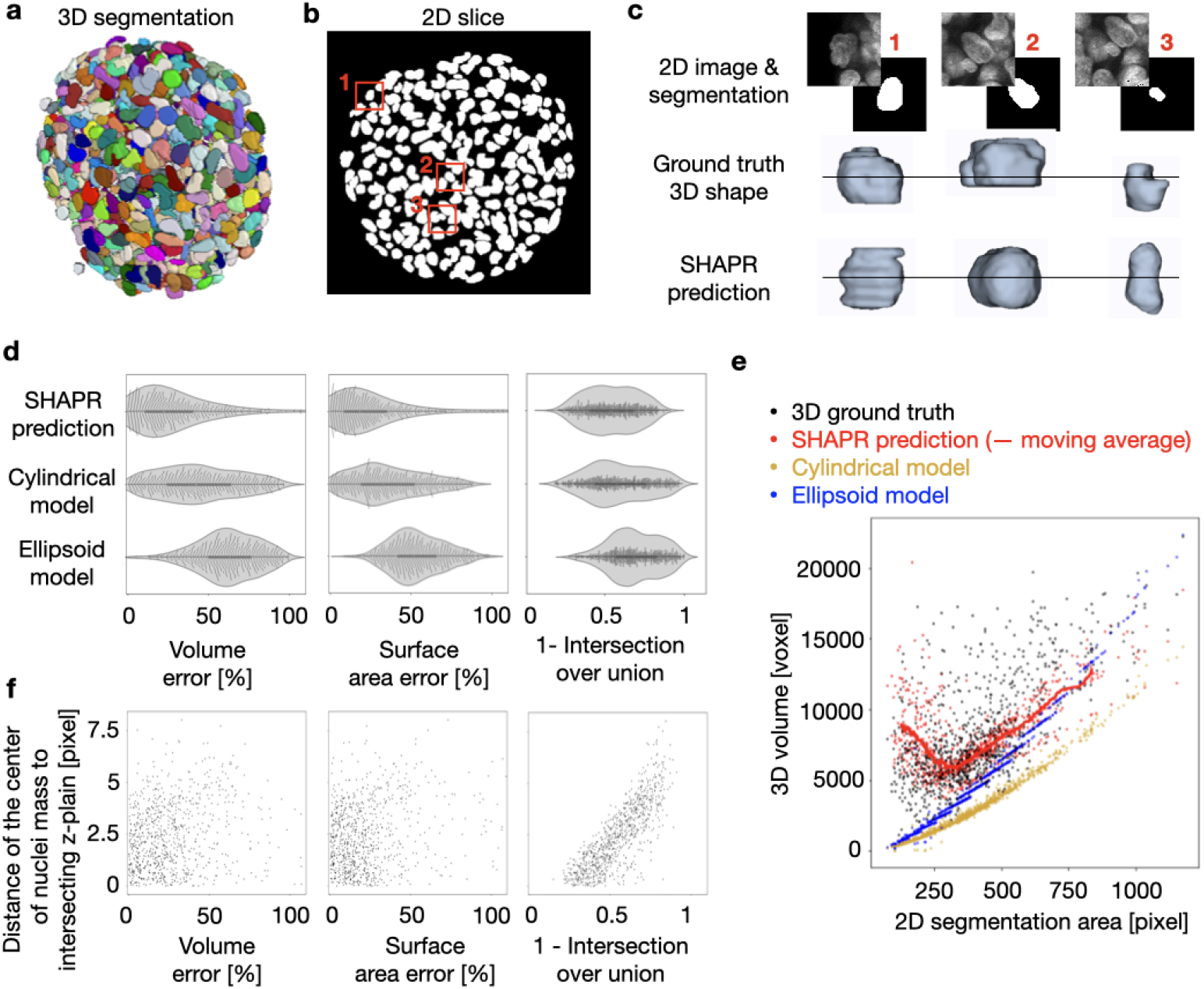
SHAPR learns fundamental 3D shape properties of human-induced pluripotent stem cell (iPSC) nuclei from a single 2D slice. **a**, Representative image of a segmented human-iPSC derived 3D cell culture with fluorescently stained nuclei. In order to generate ground truth data, six 3D cell cultures were manually segmented. **b**, 2D nuclei segmentation from a single slice at 22μm depth. **c**, 2D segmentation areas and fluorescent image intensities varied considerably with the position of the intersecting slice. **d**, SHAPR predictions outperform cylindrical and ellipsoid models in terms of volume, surface area error and intersection over union. **e**, While the cylindrical and ellipsoid models are only able to extrapolate volumes in a naïve manner, SHAPR learned complex, non-linear relationships between the 2D segmentation area and the 3D volume of a nucleus. **f**, While the intersection over union decreased with distance to the nucleus center of mass, volume, and surface predictions were unaffected.

## Discussion

SHAPR is able to solve the ambiguous inverse problem of predicting 3D shapes of single cells and nuclei from 2D images, shown on two different datasets. While Wu et al.’s (Wu et al., 2019) approach can predict the axial position of single fluorophores from a 2D image, SHAPR performs a spatial reasoning task, considering contextual information to reconstruct the shape of cells and nuclei. SHAPR is however not able to reconstruct information that is inaccessible or coming from far away from the imaged plane. While this leads to outliers with a high reconstruction error, SHAPRs was generally able to retrieve real-world 3D information and outperformed naïve 3D shape fitting models on both datasets. Furthermore we have shown that classification accuracy for red blood cells was significantly improved using the features provided by SHAPR, as opposed to the features extracted from the 2D images. Predicting 3D shapes from 2D images thus offers a simple way to reduce imaging time and data storage while retaining morphological details. A trained SHAPR model could thus enable efficient predictions of single-cell volume distributions and density, e.g. to screen organoids and identify outlier events. In combination with in-silico staining approaches (Christiansen et al., 2018; Ounkomol et al., 2018; Rivenson et al., 2019), SHAPR could be used for label-free single cell classification, e.g. for diagnostic purposes in computational pathology. We are curious to further explore SHAPR’s potential on phenotypic data, where cell morphologies may be subject to change and a model trained on wildtype data might have difficulties to generalize. As a general framework SHARP is not limited to confocal fluorescence images, and it will be particularly interesting to integrate different image modalities in the future. Alos, utilizing multiple 2D slices as an input for SHAPR could prove beneficial, as well as using more informative losses that e.g. incorporate topological information (Horn et al., 2021). Our well documented open source package of SHAPR, available on GitHub, allows for easy extension of the data loader and adaptations of the loss function used. Going beyond single-cell shape prediction, our approach may be extended to other biological structures, including organelles and proteins, and may increase the efficiency of biomedical imaging in multiple domains.

### Lead Contact

Further information and requests for resources and reagents should be directed to and will be fulfilled by the Lead Contact, Carsten Marr.

## Data availability

The datasets are available at: https://hmgubox2.helmholtz-muenchen.de/index.php/s/YAds7dA2TcxSDtr

## Code availability

SHAPR is available as a well documented, pip installable package together with commented analysis scripts at https://github.com/marrlab/SHAPR

## Methods

### SHAPR

Our SHApe PRediction algorithm SHAPR *S* consists of an encoder and a decoder with parameters θ (see Supplementary Figure 1a) that transforms an 2D input *i* ∈ *I*, which is a 2D fluorescent image and a corresponding binary mask (see Fig. 1a), to a binary 3D output *p* :

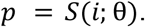

A discriminator *D* with parameters τ tries to distinguish if a 3D shape *x* comes from SHAPR or from real data:

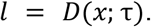

Parameters θ and τ are learned during training when the objective function ℒ is minimized:

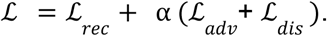

Here, ℒ_*rec*_ is the reconstruction loss, ℒ_*adv*_ is the adversarial loss, ℒ_*dis*_ is the discriminator loss and α regulates the impact of adversarial and discriminator loss during training. The reconstruction loss tries to match the generated 3D output *p* with 3D ground truth *y* and is defined as:

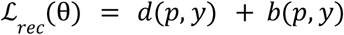

where *d*(.,.)and *b*(.,.) are Dice loss and binary cross entropy loss, respectively (see Supplementary Figure 1a). The adversarial loss tries to match the distribution of generated shapes with the dataset ground truth:

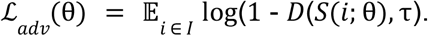

The discriminator loss is defined by:

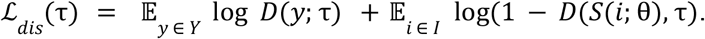

In the following implementation details are explained in further details: The encoder is built of three blocks (see Supplementary Figure 1a). Each block contains two 3D convolutional layers with a kernel size of (1,3,3), followed by a batch normalization, a dropout, and a max pooling layer to downsample convolutions. The last activation function of the encoder is a sigmoid. The decoder consists of seven convolutional blocks and each of them contains two 3D convolutional layers, followed by batch normalization, a dropout, and a 3D transpose convolutional layer for upsampling. We upsample the z-dimension seven times and the x-y dimensions 3 times in an alternating fashion. The discriminator *D* consists of five convolutional layers with a kernel size of (3,3,3) followed by an average-pooling in each dimension and two dense layers, one with 128 and one with 1 unit, followed by a sigmoid activation function, which outputs a binary label for each 3D input shape. The regularization parameter α is a step function starting with 0 so the model is trained using the reconstruction loss alone. After 30 epochs or if the validation loss has not improved for 10 epochs, α switches to 1. From then on, SHAPR is training in an adversarial fashion. Model was implemented using Tensorflow and Keras (Abadi et al., 2016; Chollet and Others, 2015).

### Training parameters

Five independent models were trained on both datasets in a round-robin fashion, so each input image was contained in the test set exactly once. For each model 20% of the dataset was used as a held out test set. 20% of the remaining data was used as a validation set during training. The remaining 60% was used to optimize SHAPRs model weights during training. SHAPRs hyperparameters, such as the learning rate and number of model weights have been fixed before training. Adam optimizer (Kingma and Ba, 2014) with an initial learning rate of 1*10^−3^, beta1 of 0.9, and beta2 of 0.999 were used. For data augmentation, training data was randomly flipped horizontally and vertically and rotated with a chance of 33% for each augmentation to be applied on each data point.

To obtain a binary image all SHAPR predictions are thresholded at 126, as their pixel values range from 0 to 255.

### Evaluation metrics

For comparison with different models, five metrics are used: relative voxel error, relative volume error, relative surface error, relative surface roughness error and intersection over union (IoU). With *Y* being the ground truth and *P* the prediction, these are defined as:

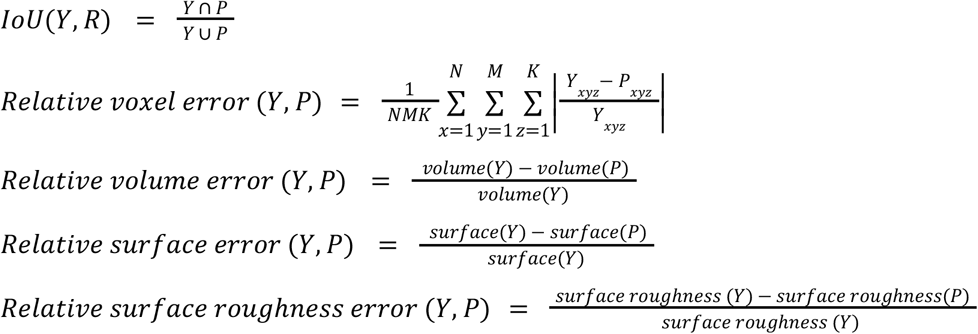

where *N, M*, and *K* are the bounding box sizes and *volume*(.) and *surface*(.) yield the volume by counting non-zero voxels:

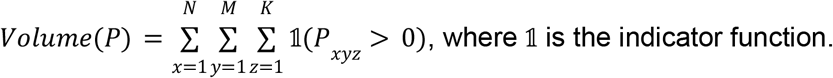

and surface area by counting all voxels on the surface of a given 3D binary shape:

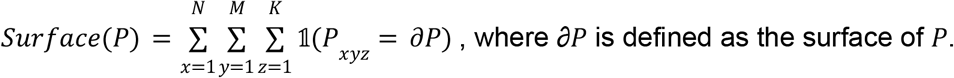

The function *surface roughness*(.) is defined as:

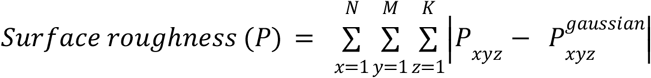

with:

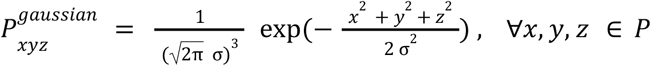

### Feature extraction

We extract 126 features from each 3D shape, comprising volume, surface, shape index, roughness, convexity, and gabor features with NumPy and the Skimage toolbox (Supplementary Table 3) (Boulogne et al., 2014). Eleven features are derived by describing the object with a mesh consisting of faces and vertices. The mesh is calculated using the marching cubes algorithm in python’s Skimage toolbox (Boulogne et al., 2014; Lewiner et al., 2003), resulting in faces and vertices. The surface, 19 mesh principals, the first nine mesh inertia eigenvalues were calculated using trimesh (“Basic Installation — trimesh 3.9.24 documentation,” n.d.). The moments of inertia represent the spatial distribution of mass in a rigid 3D shape. This depends on the 3D shapes mass, size, and shape. The moment of inertia is calculated as the angular momentum divided by the angular velocity around a principal axis. We also calculate the objects’ moments up to the third order, correlation and dissimilarity of the gray level co-occurrence matrices, and one gabor feature for each z-slice, resulting in 64 gabor features all using the Skimage toolbox (Boulogne et al., 2014).

From 2D segmentations, we extract 9 features. These are the mean pixel value, area, boundary length, boundary roughness, convexity, two moments, and three gabor features and 5 features from the 2D microscopy images (see Supplementary Table 3).

### Feature-based classification

To establish the baseline for the feature-based classification, we extract 9 morphological features (mean pixel value, area, boundary length, boundary roughness, convexity, two moments, and three gabor features) and 5 features from the 2D microscopy images that have been multiplied with the respective segmentation mask to reduce noise (mean and standard deviation of the pixel intensity, one gabor feature, and the correlation and dissimilarity of the gray level co-occurrence matrices). In total, we extract 15 features from each of the paired 2D images and 2D segmentations. We compared random forest, decision tree, K-nearest-neighbors, linear discriminant analysis, naïve Bayes, and support vector machine classifiers using the Sklearn toolbox (Varoquaux et al., 2015). The random forest classifier with 1000 estimators performed best.

Prior to the classification we oversample the training data to compensate for class imbalance, and normalize all features by subtracting their mean and dividing by their standard deviation. Prior to the tenfold cross-validation classification, we trained one random forest model to investigate feature importance. We found that accuracy increases if we remove features with an importance lower than 0.005 (see Supplementary Fig. 1c). We do not only achieve an overall higher F1 score using ShapeAEs predictions, but increase the number of true positives for four of the six classes, while for two classes we obtain the same scores (Supplementary Fig. 1a).

### Datasets

#### Red blood cells

We use 825 publicly available 3D images of red blood cells (Simionato et al., 2021) of size (64,64,64) voxels, each assigned to one of the following six classes: SDE shapes, cell cluster, multilobate, keratocyte, knizocyte, and acanthocyte. Spherocytes, stomatocytes, discocytes, and echinocytes are combined into the SDE shapes class, which is characterized by the stomatocyte–discocyte–echinocyte transformation (Chen and Boyle, 2017). The other classes’ shapes occur in samples from patients with blood disorders or other pathologies.

The number of cells in each class were: 602 for SDE shapes (93 spherocytes, 41 stomatocytes, 176 discocytes, and 292 echinocytes), 69 for cell clusters, 12 multilobates, 31 keratocytes, 23 knizocytes, and 88 acanthocytes. Red blood cells were drawn from 10 healthy donors and 10 patients with hereditary spherocytosis via finger-prick blood sampling and then fixed. Thereafter the cells were imaged with a confocal microscope and then manually classified. We extracted the 2D image from the central slide of each 3D image and segmented it by thresholding. Also, the 3D ground truth was obtained by thresholding.

#### Human pluripotent stem cells derived 3D cultures

Six human induced pluripotent stem cell-derived 3D cultures were imaged with a Zeiss LSM 880 Airyscan inverted confocal microscope with DAPI as a nuclear counterstain. Full 3D stacks were acquired using a 20X objective with a resolution of 0.25μm/pixel and a distance between slices of 1μm. We rescaled the images to 0.5μm/pixel in x-y dimension. From the center z-slice of each 3D cell culture, we manually segment and isolate all nuclei in 2D and the corresponding 3D nuclei, resulting in a dataset of 887 paired 2D/3D single-cell shapes of shapes (64,64) pixel and (64,64,64) pixel. We interpolated the single 3D nuclei to an isotropic spacing of 0.5μm/pixel. While for the red blood cell dataset we could expect to cut each single cell roughly in its middle, in this dataset the nuclei are cut at any possible z-position.

## Acknowledgments

We thank Mohammad Mirkazemi and Ron Fechnter for reviewing our code. We thank Sophia Wagner, Tingying Peng, Sayedali Shetab Boushehri, and Matthias Hehr for discussions and for contributing their ideas. We thank Marius Bäuerle and Valerio Lupperger for their feedback on the figures and manuscript.

## Author contributions

DW and NK implemented code and conducted experiments. DW, NK, and CM wrote the manuscript with AS, SA and MM. DW created figures and the main storyline with CM. SA and MM provided the 3D cell culture dataset and ideas. CM supervised the study. All authors have read and approved the manuscript.

